# Cells function as a ternary logic gate to decide migration direction under integrated chemical and fluidic cues

**DOI:** 10.1101/2022.04.28.489798

**Authors:** Hye-ran Moon, Soutick Saha, Andrew Mugler, Bumsoo Han

## Abstract

Cells sense various environmental cues and process intracellular signals to decide their migration direction in many physiological and pathological processes. Although several signaling molecules have been identified in these directed migrations, it still remains elusive how cells decipher multiple cues, specifically chemical and fluidic cues. Here, we investigated the cellular signal processing machinery by reverse-engineering directed cell migration under integrated chemical and fluidic cues. We exposed controlled chemical and fluidic cues to cells using a microfluidic platform and analyzed the extracellular coupling of the cues with respect to the cellular detection limit. Then, the cell’s migratory behavior was reverse-engineered to build the cell’s intrinsic signal processing system as a logic gate. Our primary finding is that the cellular signal processing machinery functions as a ternary logic gate to decipher integrated chemical and fluidic cues. The proposed framework of the ternary logic gate suggests a systematic approach to understand how cells decode multiple cues to make decisions in migration.

## Introduction

Directed cell migration is ubiquitous in many physiological and pathological processes, including cancer metastasis, embryonic development, inflammation, wound healing, and angiogenesis [1-6]. During these processes, cells sense and process multiple and often heterogeneous cues. These cues are chemical, mechanical, and fluidic ones [4,7-9]. Even though extensive research has been performed to identify key signaling molecules for various environmental cues, it is still puzzling how cells decipher simultaneous heterogenous cues and decide on a migration direction.

Cells can sense a chemical cue – a concentration gradient of chemokines or growth factors – through corresponding receptors on the cell surfaces, including G-protein coupled receptors (GPCR) and receptor tyrosine kinases (RTK) [5,10,11]. Cells can also sense a fluidic cue for directed migration [12-14]. Although it has not been fully understood, shear flow sensing has been considered either by force transmission through integrins for endothelial cells [15,16] or surface glycocalyx for cancer cells [17], or autologous chemotaxis involving ligand secretion and detection near the cell surface [18]. After sensing these chemical or fluidic cues, cells transduce the cues into the migratory signal via complex intracellular pathways to execute the directed migration. For instance, RTKs locally activate GTPases through the Rho subfamily, phosphoinositide3-kinase (PI3K), and ROCK/LIMK/cofilin pathways when detecting corresponding chemical cue to regulate actin polymerization, microtubule dynamics, and adhesion dynamics, eventually governing cellular polarization and asymmetric force generation for directed migration [5,19-23]. Furthermore, fluidic cue sensing can steer the directed migration by activating focal adhesion kinases (FAK) through integrin, ERK, and PI3K [15,24-26]. Indeed, cell trajectories were mostly aligned to the flow streamlines with FAK activation, where the FAK are signaling networks governing mechanotransduction involved in local activation of the Rac pathway to govern actin dynamics [24,25]. T lymphocytes could also sense the fluidic cue and showed directed migration toward the upstream direction of blood flow requiring LFA-1 of T-cell integrins and corresponding pathways such as PI3K and ERK [27,28]. Besides investigating molecular pathways of directed cell migration, the cellular sensing and processing machinery has been modeled as a biological processor in synthetic biology [29-31]. The synthetic models illustrate the cellular signal processing machinery composed of signal inputs (sensing), a logic system as a processor and an actuator (processing through complex intracellular signal networks), and outputs (cellular responses).

Despite advances in understanding the effect of either a chemical or fluidic cue alone, how cells respond to integrated chemical and fluidic cues is still not well understood. Cellular response to multiple cues has been studied in the context where both cues are chemical. In many cases, exposing cancer cells to two growth factors showed a synergistic effect on cell motility [32-35]. When one of the growth factors stimulates cells in the form of gradient, the other can have either a synergistic [36,37] or antagonistic [38] effect on directional accuracy or motility for directional migration. While the synergistic combination of the chemical cues was shown from the cooperative effect of their downstream pathways [39,40], antagonistic results were illustrated with cells’ signal-processing capacity [38]. The cell’s ability to sense and process multiple chemical cues simultaneously has been physically modeled to predict limits of the cellular ability [41,42] or to distinguish one chemical from another [43-45]. Nonetheless, in comparison to integrating multiple chemical cues, the integration of chemical and fluidic cues has been understudied, although cells are exposed to both chemical and fluidic cues *in vivo* [16,46].

In the present study, we investigated the cellular signal processing machinery by reverse-engineering directed cell migration to elucidate a biophysical understanding of how cells decipher integrated chemical and fluidic cues to determine migration direction. We exposed controlled chemical and fluidic cues on a murine pancreatic cancer cell line (KIC) in the collagen matrix using a microfluidic platform and analyzed extracellular complication of the cues with respect to cellular detection limit. Specifically, we applied pressure-driven flow to the cells that were simultaneously exposed to the TGF-β gradient in two scenarios: 1) parallel flow of an additive cue with the TGF-β gradient and 2) counter flow of a competing cue to the TGF-β gradient. Under these integrated cues, we characterized the directional accuracy of cell migration. The results were reverse engineered to construct cell’s intrinsic signal processing system as a logic gate. The results were further discussed to lay the groundwork of a systematic approach to understand how cells decode multiple cues to make decision in migration.

## Results

### Creation of a cellular microenvironment with controlled chemical and fluidic cues

To evaluate the effect of the integrated chemical and fluidic cues, we engineer the cellular microenvironment by using a microfluidic platform having a center and two side channels [38,47]. A center channel contains cells embedded in a type I collagen mixture in the platform, where two adjacent source and sink channels are filled with the medium. The chemical gradient and pressure-driven flow are simultaneously developed in the center channel by manipulating both chemical concentration and pressure variances between source and sink channels as described in **Materials and Methods**. Here, we consider two combinations based on the flow direction: parallel and counter flow (**Figure 1A)**. A parallel flow is represented as a positive direction (+) to the chemical gradient where the flow direction is from the higher to lower concentration of the chemical. On the other hand, the direction of the counter flow is represented as a negative direction (-) to the chemical gradient flowing from lower to higher concentration. By using the platform, we investigate the migration behaviors of cells under the engineered environment of integrated chemical and fluidic cues. We use murine pancreatic cancer cells (KIC cells) whose directed migration is stimulated by the TGF-β gradient [10,38,48]. The directed cell migration is often characterized by its directional accuracy, directional persistence, and motility [47,49]. In this study, we focus on the directional accuracy, representing how cells accurately follow the cue direction. In order to quantify the cellular directional accuracy to an environmental cue, we use a directional accuracy index (DAI; see Materials and methods) as defined in **Figure 1B**. Here, we note that the DAI distribution of the control is concentrated at the extremes of -1 and 1, this is an expected and well-known consequence of the cosine in its definition, as a uniform distribution of angles produce a nonuniform distribution of cosines that is more concentrated at the extremes [38,47,50,51].

**Figure 1.**
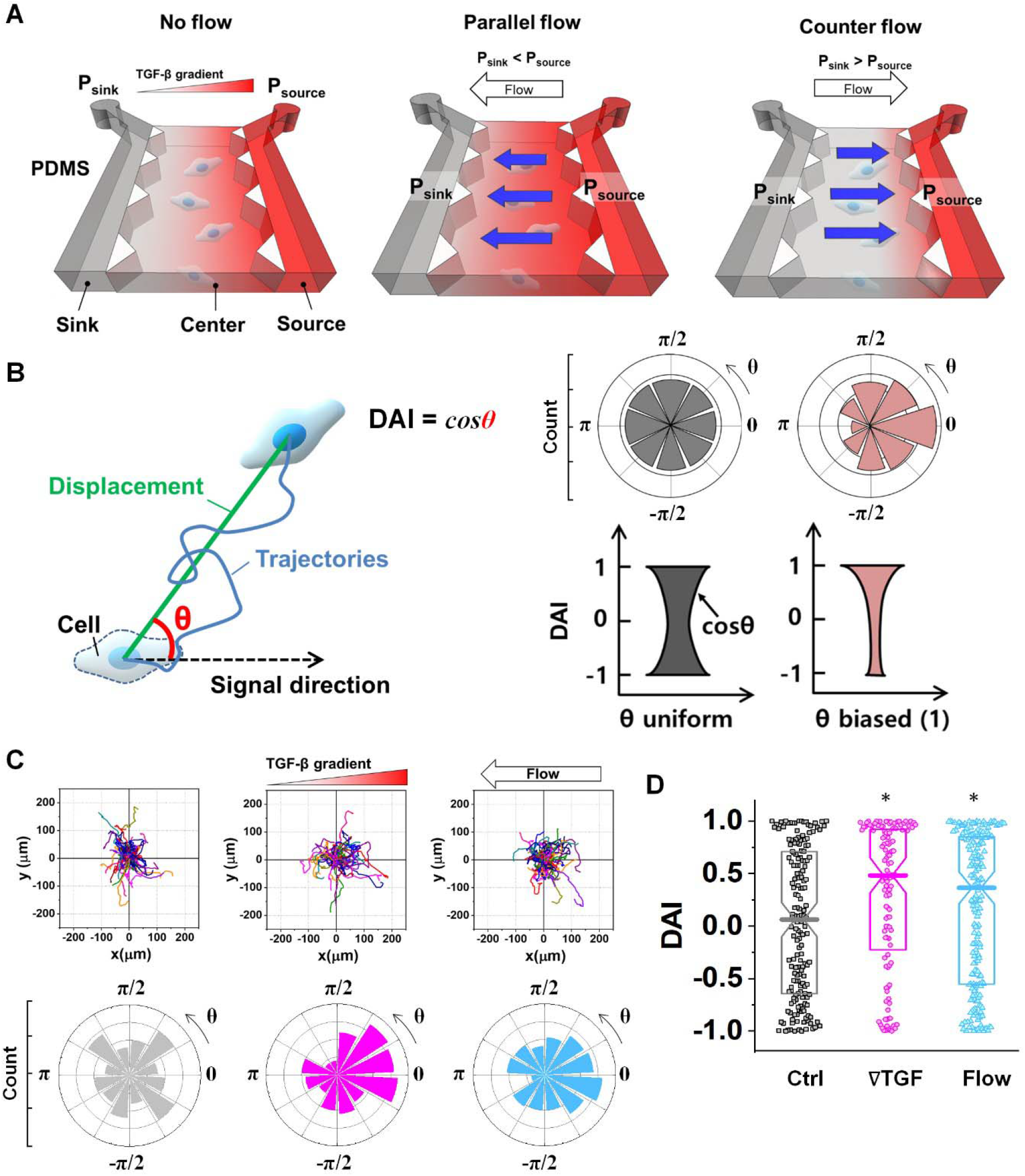
Microfluidic platform of directed cell migration under the integrated chemical and fluidic cues. (A) Schematic description of a microfluidic platform to induce the chemical gradient with pressure driven flow (Flow). Flow direction is defined based on the chemical gradient – the chemical gradient and pressure gradient is aligned; parallel flow, and the chemical gradient and pressure gradient is in opposing directions; counter flow. (B) Directional migration is characterized with directional accuracy index (DAI) defined as a cosine of the angle (θ) between the cue and displacement direction. (C) Representative cell migration trajectories of control (Ctrl, grey), 10nM/mm TGF-β gradient (∇T, magenta), and interstitial flow (Flow, cyan) and angular distribution for θ respectively (D) DAI distribution of collected cell trajectories of Ctrl, ∇T, and Flow. Box: quartiles with a median line in the middle of the box. Dot: the corresponding metric from a single trajectory. *: p < 0.05 (Mann-Whitney U-test)

As a result, the KIC cells showed significantly enhanced directional accuracy responding to the TGF-β gradient (**Figure 1C, magenta)**, consistent with previous studies [38,48]. The directed migration is notably induced by a TGF-β gradient so that the DAI is biased towards 1, indicating that the cells’ movement is biased toward a (+) direction (**Figure 1D, magenta**). In contrast, the control group is unbiased as a median of the DAI distribution is close to 0 (**Figure 1D, gray**). In addition to chemical cues, we observed the flow-induced directed migration of KIC cells, as shown in **Figure 1C** (**cyan**), when the cells were exposed to the flow of approximately 1.5 μm/s (**See details in Materials and Methods section and Figure S1**). The DAI distribution of cells in response to the flow is biased toward 1, indicating that the cells move to the upstream flow direction as reported previously [25]. The DAI distribution of cells in response to the flow is significantly biased compared to the control, as shown in **Figure 1D**. These results confirm that the KIC cells respond to chemical and fluidic cues.

### Extracellular combination of the chemical and fluidic cues creates regions where the chemical cue becomes below the cellular sensing limit

Chemical cues in the cellular microenvironment are transported by not only diffusion but also interstitial fluid flow [52-54]. To characterize this extra-cellular complication, the concentration profiles of a chemical cue in the presence of the flow on the microfluidic platform were measured and predicted by using FITC-conjugated dextran in **Figure 2**. The intensity measurement was considered as concentration of the FITC-dextran. Without flow, the concentration gradient of a chemical cue is a linear profile (**Figure 2A**). When the interstitial fluid flow of 1.5μm/s was imposed along the chemical cue (i.e., parallel flow configuration in **Figure 2B**), the gradient becomes shallow in the region of interest (ROI) except the edge region (*x* ∼ 250 um). Since the parallel flow augments the advection of the molecules along the chemical cue gradient, the overall concentration value increases (**Figure 2B**). On the contrary, the counter flow suppresses the chemical cue gradient and lower the overall chemical cue concentration. Near the edge of source side (*x* ∼ 750 um), the gradient grows and becomes steep (**Figure 2C**). This result demonstrates that the concentration gradient of chemical cues in the microenvironment is significantly altered by the presence of the interstitial flow. Considering the interstitial flow can also regulate the directed cell migration as a fluidic cue, cells under chemical and fluidic cues need to process much more complex extra- and intra-cellular signals.

**Figure 2.**
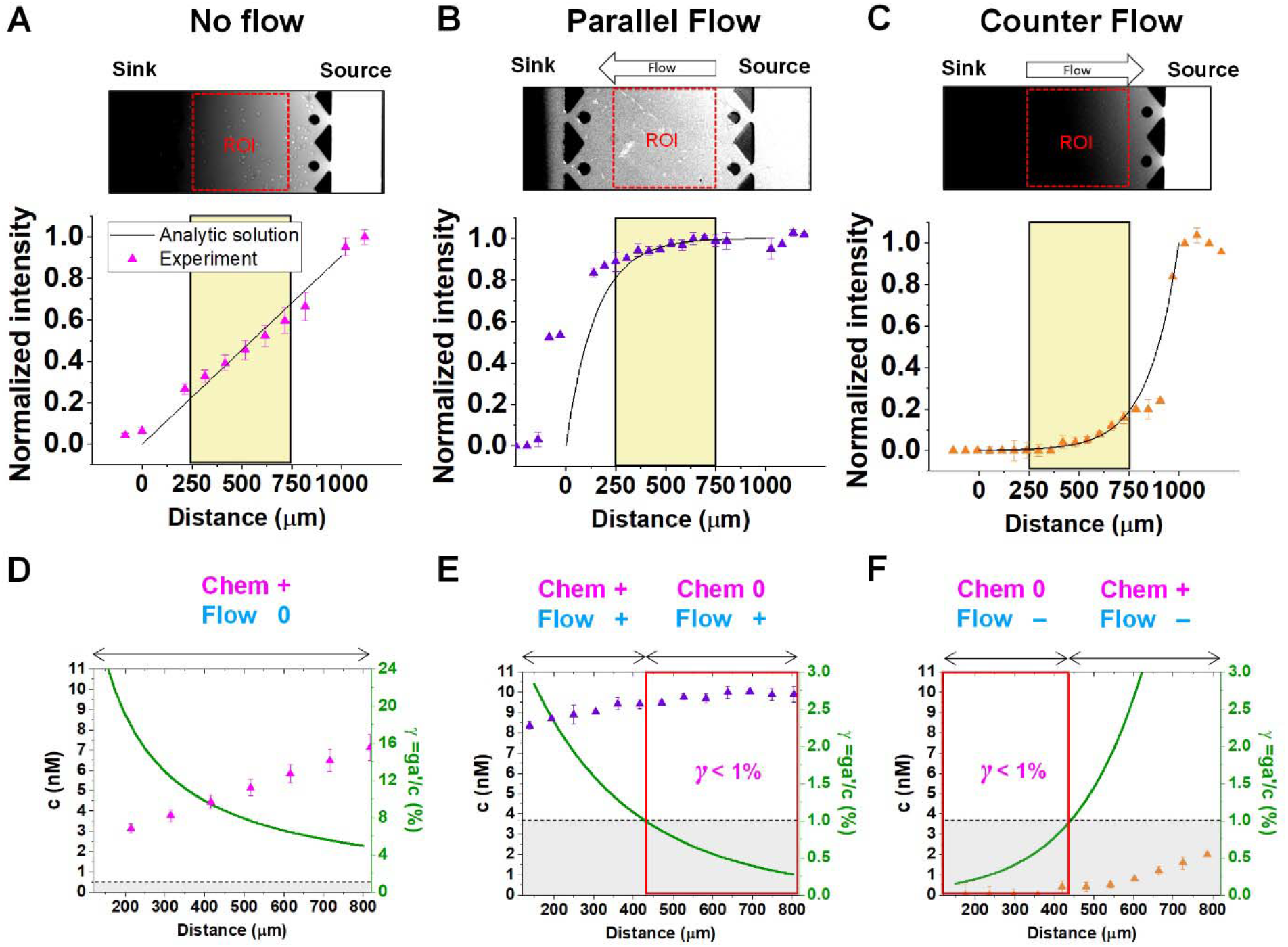
Extracellular complications of the integrated chemical and fluidic cues with cellular detection limit. Concentration profiles of a chemical cue with (A) no flow (magenta), (B) parallel flow (purple), and (C) counter flow (orange) is simulated by 10kDa FITC-dextran. Concentration data points were measured from fluorescence intensity of FITC-dextran across y-axis (mean ± S.D.). Solid lines represent analytic prediction. The yellow region indicates Region of Interest (ROI) where cell trajectories are analyzed excluding any edge effect of the microfluidic platform. A relative gradient of the chemical concentration across the cell body (*γ*, green) was calculated based on the corresponding concentration profiles of (D) no flow, (E) parallel flow, and (F) counter flow of ROI. The signal state of the chemical cue (Chem, magenta) was defined as detectable when *γ* >1% whereas not detectable when *γ* <1%. The fluidic cue is represented as Flow (dark cyan). A dot represents mean ±S.D. Red box represents *0-state* indicating that negligibly shallow gradient which cells are not capable of sensing.

Then, we analyzed the complication of the integrated chemical and fluidic cues asking if the non-linear cue profiles fulfill the physical detection limit for chemical cue. The physical detection limit for chemical cue is a cellular capacity physically governed for a shallow chemical gradient [55,56]. Although the exponential profiles (either parallel or counter flow) provided a steep gradient near the source or sink, most of cells were located in the area where a relatively shallow gradient is present. The physical detection limit was roughly determined with a relative gradient of the chemical concentration across the cell body (*γ*) as follows [47,55,57] :

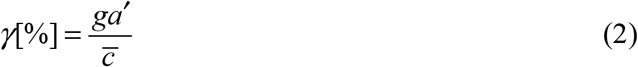

where g [nM/mm] indicates a gradient strength, *a*’ is the estimated cell length, and 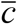 is an average concentration (See Materials and Methods and **Figure S2**). We determined the cellular detection precision with *γ* ∼ 1% as a physical detection limit, as the cells may not be capable of sensing the chemical gradient below this limit based on knowledge of the sensory precision threshold for *Dictyostelium* [58,59] and cancer cells [18,47]. Here, we defined the cue directions as forward (+ state), backward (– state), and no-cue (0-state). If a gradient is present but below the detection limit for the cells (*γ* < 1%), the gradient is neglected by the cells. Consequently, it is also considered as a *0-state*, indicating that there is no gradient which cells can sense.

In the no-flow condition (**Figure 2D**), all regions were above the physical detection limit, indicating that the cells are capable of sensing the chemical gradient. On the other hand, both parallel and counter flow conditions presented in **Figure 2E and F** display ‘*0-state*’ regions where the relative gradient is below the cells’ physical detection limit, leading to differential signal environment in two ways. For the parallel flow, the *γ* value drops down as the location is close to source channel and gets to the detection limit (*γ* ∼ 1%) in the middle of ROI shown in **Figure 2E**. Consequently, it divides the region into two where the chemical cue is detectable (chem + state) and not detectable (chem *0-state*). When the chemical cue is detectable, cells are exposed by additive combination of the chemical gradient and the flow. Interestingly, *0-state* in parallel flow, the background concentration of chemoattractant is close to 10nM. We anticipate that it is equivalent to the situation of cells exposed to a uniform chemoattractant with flow. On the other hand, *γ* for the counter flow increases as it is close to source channel while the detection limit (*γ* ∼ 1%) is in the middle of ROI (**Figure 2F)**. At the location where the chemical cue is detectable, the combination of the chemical and fluidic cues is competitive, having opposite direction (Chem +/Flow – state). Unlike the parallel flow, the counter flow washes the chemoattractant mostly away from the ROI showing the background concentration as close to 0nM where the chemical cue is below the detection limit in **Figure 2F**.

### Intra-cellular processing of two cues simultaneously

**Figure 3**. shows the directed migration behaviors of KIC cells under integrated chemical and fluidic cues. The cells’ migration trajectories and the angular distribution of corresponding displacement are presented in **Figure 3A and B**. The results are divided into sub-regions considering the signal states of (TGF-β gradient/flow). For parallel flow (**Figure 3A**), the trajectories and angle (θ) are distributed biased features toward the chemical cue direction in the sub-region of the additively integrated chemical and fluidic cues (+/+), whereas the trajectories and their angles in the other sub-region of 0/+ are randomly distributed. For the counter flow presented in **Figure 3B**, the trajectories and angular distribution are biased toward the chemical cue direction in the sub-region of the competitively integrated cues (+/–) whereas those are biased toward the flow direction in the other sub-region of 0/–.

Resulting directional accuracy is further analyzed with directional accuracy index (DAI) of all experimental cases in **Figure 3C**. DAI of the cell trajectories under a single cue either TGF-β gradient (**Figure3C, magenta**) or flow (**Figure 3C**, dark cyan) are biased toward each cue direction whereas control DAIs (**Figure3C**, grey) show a distribution with median close to 0. In the parallel flow, the directional accuracy is significantly enhanced toward the chemical cue direction in +/+ as shown in **Figure 3C (purple, left)**. Indeed, the DAI distribution is highly biased toward 1, with a median as 0.62 in this case. Although the gradient strength is ∼10% shallower than a linear TGF-β gradient, directional accuracy under +/+ state is still significantly biased toward the cue direction comparable to the linear gradient (median DAI=0.46). On the other hand, cells lose their directional accuracy completely under 0/+ state where *0-state* for the TGF-β gradient despite the flow presence shown in **Figure 3C (purple, right**).

**Figure 3.**
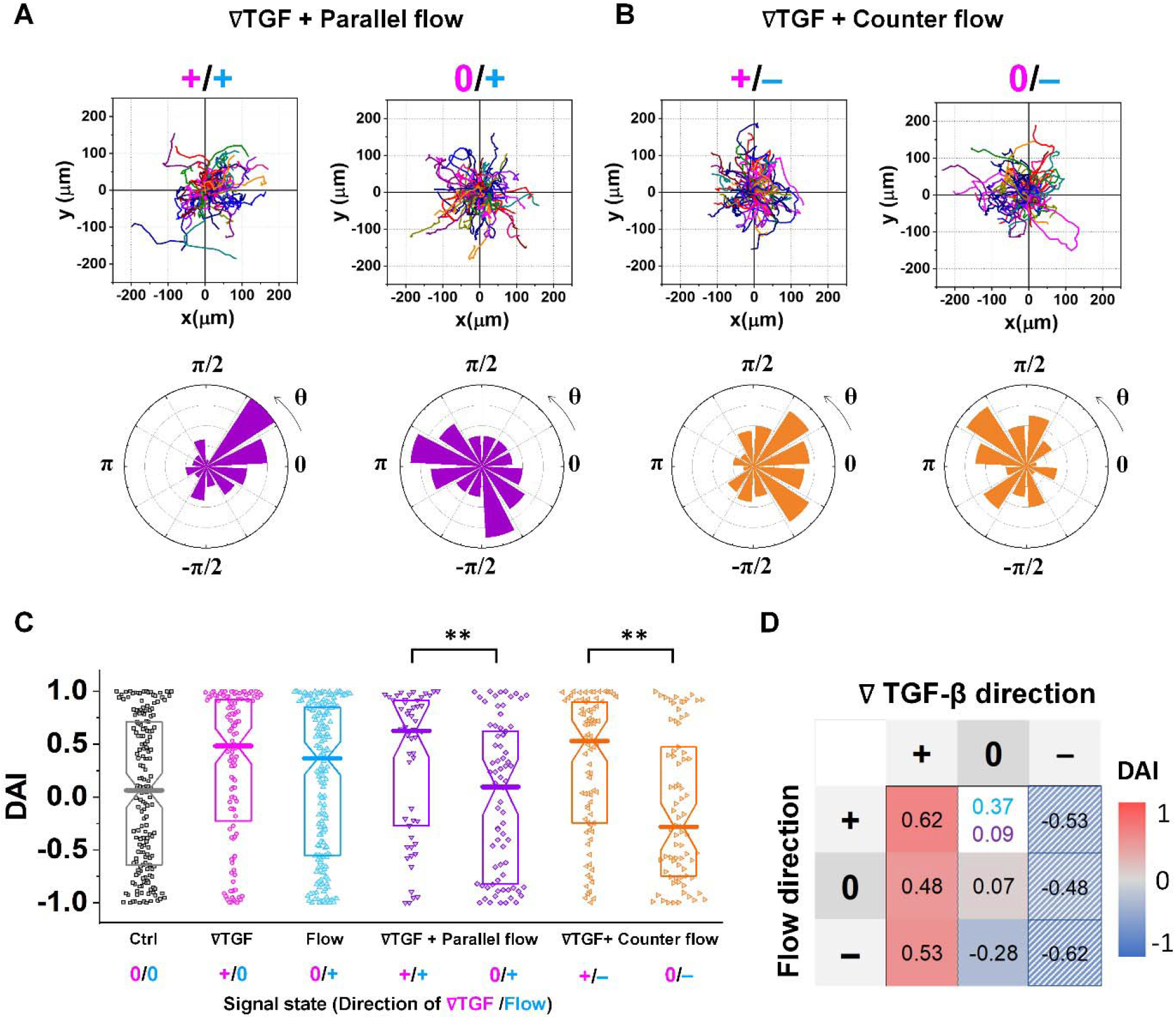
Differential response in directional accuracy of KIC to the integrated cue. Cell migration trajectories and angular distribution (θ) of collected trajectories of KICs under (A) TGF-β gradient (C_source_=10nM and C_sink_=0nM) with the parallel flow (∇TGF + Parallel flow, purple), and (B) TGF-β gradient with the counter flow (∇TGF + Counter flow, orange). (C) DAI distributions of all collected trajectories of KICs with respect to each signal state of ∇TGF (magenta) / Flow direction (dark cyan). Box: quartiles with a median line in the middle of the box. Dot; a DAI from a single trajectory.; Cell trajectories N>50. **: p<.01, (Mann-Whitney test) (D) Heat map for medians of DAI distributions of all experimental conditions. The hatched area: reflected from the opposite signal state.

The TGF-β gradient and the counter flow compete in their directions when stimulating the cells. Here, we define reference direction for DAI as TGF-β gradient direction, resulting in a negative sign for the directed migration stimulated by the flow. Cells under the counter flow in the region with the TGF-β gradient above the limit (+/–) show bias in their DAI distribution toward 1, showing a median DAI = 0.53 (**Figure 3C, orange, left**). Although the counter flow direction is the opposite of the TGF-β gradient, cells remain significantly biased toward the TGF-β gradient. On the other hand, cells under the counter flow with *0-state* of TGF-β gradient (0/–) have biased distribution of DAI toward -1 with a median as -0.28 (**Figure 3C, orange, right**). It implies that cells are not capable of sensing the shallow chemical gradient in the 0-state region, consequently, they respond only to the flow.

We summarize the median DAI from distributions of each signal state in the heat map (**Figure 3D**) to show how each signal state induces the directional accuracy. The signal states with negative chemical cue direction (–/+, –/0, and –/–) are simply reflected by the signal states (+/–, and +/0, and +/+, respectively). The heat map shows two distinct features. Regardless of the fluidic cue, cells seem to follow the chemical cue direction when the chemical cue is not *0-state*. Indeed, the cells seem to neglect the flow when they are exposed to a competing combination of TGF-β gradient and the counter flow. If the cellular response simply follows the signal state hypothesizing that the chemical and fluidic cues have comparable level in cellular processing machinery, the signal state of (+/–) would be anticipated as an antagonism showing lower DAI than TGF-β gradient only, but this was not shown in our results. Also, the 0/+ state can be represented in two distinct ways: flow only (dark cyan) and 0/+ state of chemical cue with parallel flow under the integrated chemical and fluidic cues (purple) (**Figure 3D**). The median DAI under the flow only was 0.37, which was significantly biased toward the upstream direction of the flow. However, cells under for 0/+ of the integrated chemical and fluidic cues lose their bias completely with the median DAI = 0.09, indicating the cells do not respond to the flow stimulation. Unlike cells in 0/+, the cells in *0-state* with counter flow (0/–) were induced by the flow. Thus, a quantitative comparison of effectiveness between chemical and the fluidic cues is required to address the results. Since the *0-state* with parallel flow includes the high background concentration of TGF-β, we hypothesize that cells are receiving strong information about an ungraded chemical cue, and this overpowers the weaker fluidic cue.

### A shared pathway model successfully predicts the cellular response to integrated cues

To further understand the cell’s integrated response to both flow and chemical cues, we turn to mathematical modeling. We adapt a model that we previously introduced to describe a cell’s integrated response to two chemical signals [38] that relies on the convergence of the two response pathways at a common intracellular component. Specifically, here we suppose that TGF-β induces the production an internal chemical species X, whereas flow induces (e.g., via pressure-sensitive receptors) the production of a second internal species Y (**Figure 4A**). X and Y converge to jointly catalyze the conversion of a third species A into an activated state B, which is responsible for initiating the migration machinery downstream, described in the model as species M. The net result is that a rightward TGF-β gradient, or a leftward flow (corresponding to a rightward pressure gradient), produces more M molecules on the right side than on the left side of the cell, inducing rightward migration. Simplifying the cell to just these two halves, the rate equations corresponding to the reaction network in **Figure 4A** give a steady-state molecule number difference of (see **Supplementary Information**)

**Figure 4.**
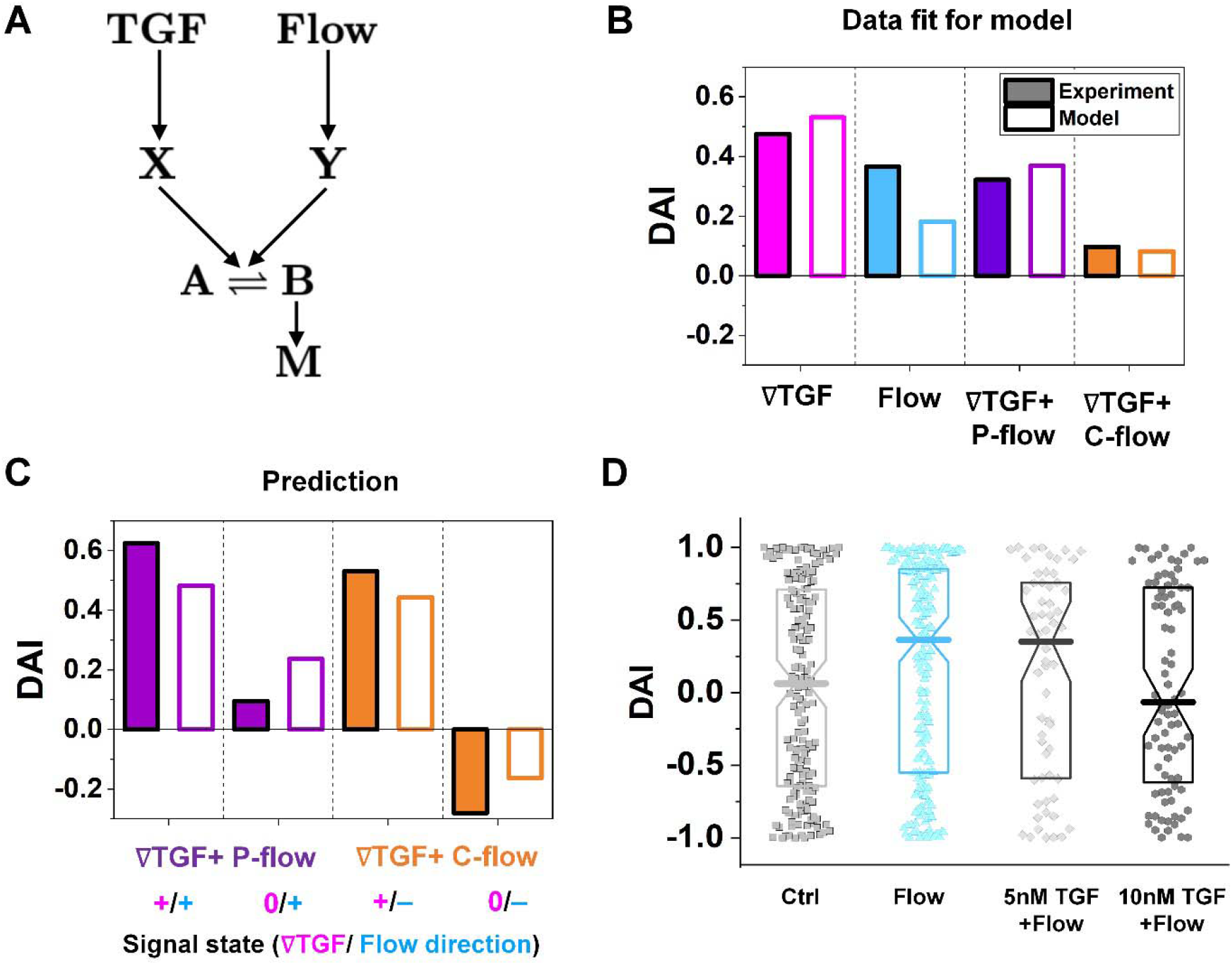
The shared pathway model addressing experiment findings under the integrated chemical and fluidic cues. (A) Simple molecular network used to explain the experimental data. (B) Fit of the experimental data using our model. (C) Prediction by our model and validation by experiments. (D) DAI distribution of KIC cells migrating in response to flow and background TGF-β present together. Box: quartiles with a median line in the middle of the box. Dot; a DAI from a single trajectory.; Cell trajectories N>50.

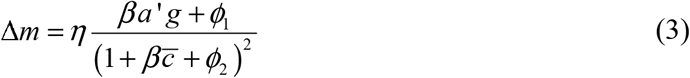

where, as above, 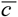 is the background TGF-β concentration in the region of interest, _*g*_ is its gradient, and *a*’ is the cell length; and here *η, β, ϕ*_1_, and *ϕ*_2_ are combinations of reaction rates (see **Supplementary Information**). Intuitively, *η* sets the overall molecule number scale, *β* is an amplification factor for the chemical signal, and *ϕ*_1_ and *ϕ*_2_ depend on the properties of the flow.

To describe the resulting migration, we use a biased random walk model [47] to relate the migration angle *θ* to the molecule number difference Δ*m*,

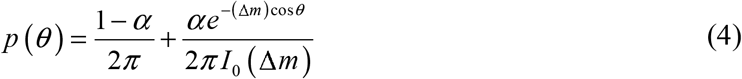

Here *p* (*θ*) is the probability distribution of migration angles (**Figure 1B**), the first term corresponds to purely random motion over the angular range 0 to 2*π*, and the second term corresponds to directed migration toward *θ* =0. Intuitively, as Δ*m* increases, the second term becomes more sharply peaked, corresponding to higher directional precision. The parameter *α* determines the balance between the random (*α* = 0) and directed (*α* =1) components, and *I*_0_ is the modified Bessel function of the first kind (required for normalization).

The median of cos *θ* values drawn from *p* (*θ*) gives the DAI from the model in terms of the parameters 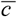, _*g*_, *a*’, *η, β, ϕ*_1_, and *ϕ*_2_ and *α*. We compare the model with the experiments in two steps. First, we calibrate the model parameters using the experimental data. Specifically, we set 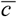, _*g*_, and *a*’ directly from the experiments as above; we set the four parameters *η, β, ϕ*_1_, and *ϕ*_2_ using the median DAI in the four experimental conditions (TGF-β gradient only, flow only, parallel flow, and counter flow); and we set the last parameter *α* using the maximum mean DAI observed across all of these experimental conditions (see **Supplementary Information**). We see in **Figure 4B** that the model is able to capture the median DAI from experiments well. Second, we use the calibrated model parameters, with no further fitting, to predict the median DAI when the parallel and counter flow conditions are separated based on the detection limit as above. We see in **Figure 4C** that the model prediction agrees well with the observed median DAI values, even without further fitting.

Beyond validating the experiments, the model offers an intuitive explanation for the cell responses. When the TGF-β and flow signals are coherent (parallel flow), and above the TGF-β gradient detection limit, the DAI is large, as expected (**Figure 4C, left purple**). Below the detection limit, one might expect that flow should dominate, and the DAI would still be positive. However, the large TGF-β background concentration in this regime (**Figure 2E** and large 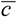 in **Eq. 3**) saturates the signaling network, leading to a small Δ*m* and thus a small DAI (**Figure 4C, purple right**). When the TGF-β and flow cues are incoherent (counter flow), and above the TGF-β gradient detection limit, the DAI is large and positive (**Figure 4C, orange left**), indicating that chemical detection overpowers flow detection. Indeed, in the model we find that *ϕ*_1_ / *ϕ*_2_, which is the analog of 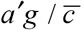 for flow sensing (**see Supplementary Information**) is 0.1%, which always less than 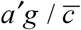 in regimes where it is above its detection limit of 1%. Finally, below the chemical detection limit, the DAI is negative (**Figure 4C, orange right**), i.e., aligned with the flow, because here the TGF-β background concentration is negligible, allowing flow to dominate.

To confirm a key prediction of the model, namely that the large TGF-β background concentration is responsible for the suppression of flow sensing in the parallel flow regime below the chemical detection limit (**Figure 3C, right purple**), we perform further experiments. Specifically, we combine flow with a uniform TGF-β concentration at either 5 or 10nM. At 5nM, which is roughly half of the background level in this regime (**Figure 2E**), we see that the DAI is not suppressed (**Figure 4D**). However, at 10nM, which is roughly equal to the background level in this regime, we see that the DAI is indeed suppressed (**Figure 4D**).

### Cellular signal processing machinery can be modeled as a ternary logic gate

We construct a logic gate model to reconstitute the function of the cellular signal processing machinery (**Figure 5**). The cellular response to the cues (+, 0, or –) presents three variables as outputs, allowing us to develop a ternary logic system. For consistency, we define the output direction based on the chemical cue. When the cell migration direction is aligned to the chemical cue direction with positive DAIs, cell direction can be represented as a forward (*+ state*). On the other hand, the repulsive response to the cue with negative DAI can be denoted as *– state*. Cells’ random movement not showing any bias in their direction with DAI close to 0 are defined as *0-state*. In this way, the heat map presented in **Figure 3D** can be converted to a ternary logic table. We convert the positive or negative DAIs to + or – respectively when the DAI distribution fulfills the statistical significance (p<0.05) in their comparison with control (**Figure 5A**). The DAIs close to 0 with no significant bias in their distribution is converted to 0. We present two separate ternary logic tables based on the system saturation caused by the high background TGF-β concentration resulting in suppression of the directional accuracy as we presented in the prior section. By separating the results depending on the system saturation, the present inconsistency presented in 0/+ (**Figure 3D**) is resolved.

**Figure 5.**
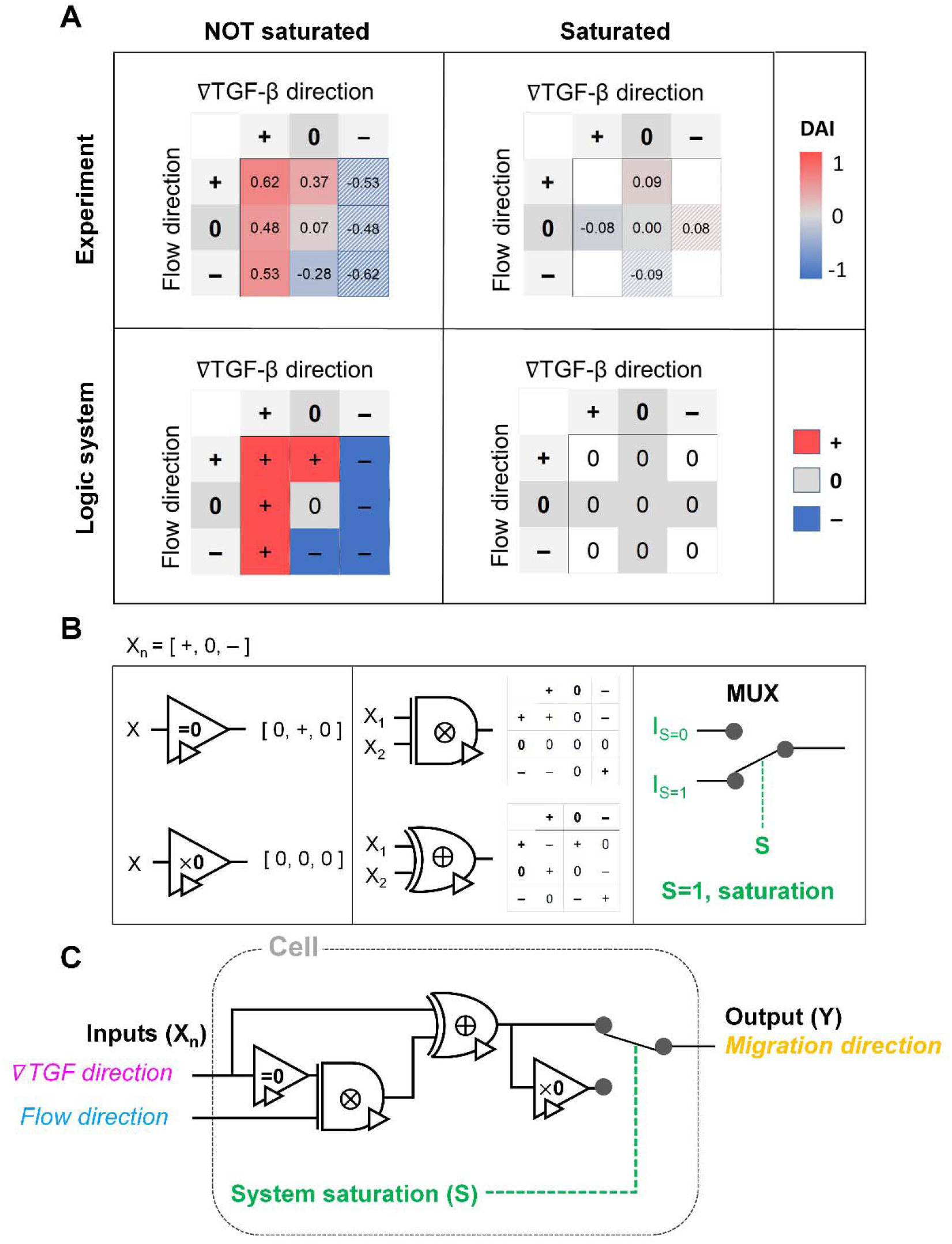
Ternary logic gate model to address the cellular signal processing machinery. (A) Heat map for experimental results of DAI medians of not saturated (left) and saturated (right) cases. The system saturation is considered with experimental groups of the higher TGF-β background noise (TGF=10nM). It is converted to the truth tables of ternary logic system with signal states (+, 0, and –); the hatched area: reflected from the opposite signal state. (B) Ternary operators and their functions used in the model. (C) The proposed ternary logic gate model.

The ternary logic gate is composed of the five ternary operators, whose operating functions are presented in **Figure 5B**. The monadic operator “=0” returns 0 input to + output whereas + and – input to 0 output. Another monadic operator “×0” returns all zero regardless of the input states. We also used diadic operators represented as ⨂ and ⨁, which simply multiply and add two inputs to return the corresponding outputs respectively. To stop misguided migration when the machinery capability is saturated, we apply a circuit breaker for the system saturation with a multiplexer. The multiplexer switches the circuit path based on an additional intracellular input *S*. We apply two intracellular inputs; S=1 where the system is saturated by high background TGF-β concentration, and S=0 where the system is not saturated. By using the basic operators, the ternary logic circuit to address the ternary logic tables is developed in **Figure 5C**. When the system is not saturated (S=0), the cells tend to decide their direction dominantly following TGF-β gradient, regardless of the flow’s existence. The path for S=0 mimics an absorption logic gate which selectively choose one particular input to decide their output. In contrast, the path for S=1 for system saturation leads to returning all zero. Consequently, the circuit successfully represents the experimental results. The ternary logic gate in **Figure 5C** implies a corresponding mathematical expression in terms of ternary variables (–, 0, or +) which, self-consistently, agrees with our expression for Δm (**Eq. 3**) when looking only at its sign (–, 0, or +); see **Figure S3**.

## Discussion

The present results show the complexity in the extracellular signal environment, specifically caused by the integrated chemical and fluidic cues. We investigated signaling environment where Péclet number (Pe) ∼ 1 in cases that flow runs parallel or counter to a TGF-β gradient. As the fluidic cue becomes stronger (i.e, a higher Pe environment (Pe >> 1)), the transport of TGF-β becomes convection-dominant, whereas weaker fluidic cues (i.e., lower Pe (Pe << 1)) corresponds to diffusion-dominant transport. Corresponding changes of the gradient of chemical cue depending on the flow direction and Pe are shown in **Figure S4**. In fact, Pe varies from 0.1 to 2 with slow interstitial flow rates in various tissue interstitium, including cancer [52-54]. The combination of the TGF-β gradient and the flow displays two important aspects. First, the TGF-β concentration profiles are non-linear exponentials, where the cells experience spatially differential gradient strengths, including a shallow gradient region close to the cellular sensing limit. The exponential profiles of the concentration could be either shallow or steep where the background concentration could be higher or lower depending on the direction of flow and chemical gradient, causing a spatially differential response of cells [47,57,58]. Second, cells are exposed to integrated cues of the chemical gradient and the flow as either additive or competitive depending on the flow direction, increasing the complexity of the cellular sensing and processing machinery both intrinsically and extrinsically.

The present results streamline the complexity by implicating cellular sensing capability for the chemical cue. The spatially varied gradient is developed by imposing convection in the microenvironment, including shallow gradient regions below the cellular detection limit [55]. Indeed, the physical limit of cells in sensing chemical gradient allowed us to decouple the integrated chemical and fluidic cues into the fluidic cue only, indicating 0-state. Consequently, the cells ruled out the effect of the TGF-β gradient in their decision-making for migration direction where it was below the detection limit.

Besides, we demonstrate the cellular response to the combination of chemical and fluidic cues. The flow impacts the cellular behaviors as a transport medium and as a fluidic cue to induce migration potential of various cell types, including immune cells and cancer [13,60,61]. In the presented experiment results, we have observed that cells effectively select a cue to follow in processing the mixed chemical and fluidic cues. When cells are capable of sensing both chemical and fluidic cues, cells tend to follow a chemical gradient direction in both the additive combination with the parallel flow and the competing with the counter flow, as shown in **Figure 3**. The cells were biased toward the upstream direction of the fluidic cue, only when the chemical gradient was too shallow for cells to detect it. (**Figure 3 orange right**). The effect of the chemical gradient is ruled out. Most strikingly, the cellular biased response was completely ruled out when the processing capacity is saturated. Based on the experimental observation, we propose the framework of the cellular sensing machinery by using the ternary logic gate model in **Figure 5**.

The present results demonstrated the physical implication of cells’ innate capability of processing the integrated cues. The cellular sensory machinery incorporates that the complex signal transduction manipulates the cellular functions after sensing the cues. Previously, we have shown that saturation of the intracellular signal transduction capacity causes antagonism in their chemotaxis, where the two different chemical cues not sharing their receptors induce cell directed migration [38]. A recent study also demonstrated that the limited source of intracellular translational or transcriptional factors results in poor performance and predictability in synthetic biology [62]. Although it is still poorly understood how flow activates cell mechanotransduction, recent studies have begun exploring the signaling cascades of the flow cue [4,14,19]. Interestingly, the downstream networks of the flow cue overlap with the chemotaxis signaling transduction that regulates actin cytoskeletal dynamics, which are thought to manipulate the cell bias movement [14]. In this sense, our results demonstrate that the saturation of the shared pathway to manipulate cellular migration direction completely removes cells’ bias movement, indicating that the cellular processing capacity could limit the cellular performance.

The present study laid a framework for understanding how cells decode chemical and fluidic cues to determine migration direction by proposing a ternary gate circuit. Cellular decision-making is a systematic result from sensing to deciphering the cues with complex downstream signal processing. Our results suggest a simple circuit to address the complex process based on our observation showing the cellular innate sensing and processing capacity [38]. The proposed framework of the gate circuit implies the potential use of the ternary system to model cellular sensory machinery for environmental cues with heterogeneous origins. The proposed ternary logic gate may provide a blueprint to synthesize functional signal processing machinery for engineered cells. Recent advances in synthetic biology to engineer genetic circuits of the cells offer great potential in developing engineered cellular systems as sensors, therapeutics, and delivery vehicles [31,63-65]. The microbials (e.g., Escherichia coli and virus) have been engineered to target pathogenic sites for diagnosis and therapeutics [66,67]. Recent development in synthetic mammalian cells pursued the immune cell (T-cell) chemotaxis [30] and anti-cancer targeting purposes [68,69]. Nonetheless, it is required to have an effective genetic circuit design to regulate the directed migration of the delivery vehicles based on a profound understanding of cellular sensory machinery with both extrinsic and intrinsic considerations. Accordingly, the proposed ternary gate model provides insight to develop potential targeting vehicles in various ways.

## Limitations of the study

Although the present study demonstrates how cells decipher integrated chemical and fluidic cues, the type of environmental cues for the investigation is limited. Multiple chemoattractants may induce directed cell migration besides TGF-β. Besides the chemical or fluidic cues, mechanical cues such as matrix stiffness gradient can also affect migration. The present study used one cell type, but further validation using multiple cell types is warranted.

## Acknowledgments

This work was partially supported by grants from the National Institutes of Health (U01 HL143403, R01 CA254110, R61 HL 159948, and P30 CA023168) and National Science Foundation (MCB-2134603, MCB-1936761, and PHY-1945018).

## Author contributions

BH conceived the idea. HM primarily performed the research and acquired the data. SS and AM performed the research and acquired the data for the shared pathway model. All authors discussed the results.

## Inclusion and diversity

We worked to ensure diversity in experimental samples through the selection of the cell lines. The author list of this paper includes contributors from the location where the research was conducted who participated in the data collection, design, analysis, and/or interpretation of the work.

## Materials and methods

### Cell cultures and reagents

KIC is a murine pancreatic cancer cell line isolated from genetically engineered mouse model for pancreatic adenocarcinoma in which *Kras* was combined with deletion of the *Ink4a* locus (*Ink4a/Arf*^*L/L*^). [70-72] The KIC cells showed mesenchymal phenotype in response to TGF-β, whose invasion potential increased and directed migration was induced [38,48]. These cells were cultured in RPMI 1640 with 2.05mM L-glutamine (GE Healthcare Bio-Sciences Corp., MA, USA) supplemented by 5% v/v fetal bovine serum (FBS) and 100 μg ml^−1^ penicillin/streptomycin (P/S). The cells were regularly harvested by 0.05% trypsin and 0.53mM EDTA (Life technologies, CA, USA) when grown to ∼80% confluency in 25 cm^2^ T-flasks and incubated at 37°C with 5% CO_2_. Harvested cells were used for experiments, or sub-cultured while maintaining them below 15^th^ passage.

### Convection-driven signal environment in a microfluidic platform

In this study, we use the *in vitro* microfluidic platform to engineer microenvironment involving both chemical and pressure variances. The *in vitro* microfluidic device is composed of center, source, and sink channels [38,47]. We manipulate concentration of transforming growth factor beta-1 (TGF-β, Invitrogen, CA, USA) between source and sink channels to develop chemical gradients in the center channel. Meanwhile, we engineer the pressure variance between source and sink channels so that the pressure driven flow is generated in the center channel. The concentration profile in the center channel could be determined by its diffusion and advection shown in the governing equation (**Eq.1**). To apply the interstitial flow in a presence of TGF-β gradient, we always filled the source channel with 10nM of TGF-β while the sink channel was filled with normal culture medium. The concentration profile of TGF-β was analyzed with simple mathematical approach through the governing equation (Eq.1) and corresponding boundary conditions, providing structural intuition of the gradient features. We simplified the device geometry as a 1-D, used constant parameters of diffusivity (*D*_*eff*_) and flow velocity (*V*_*f*_ = *U*), and evaluated the steady state 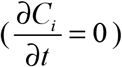.

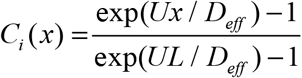

Consequently, the concentration is an exponential profile. Exponential non-linear gradient profiles are expected to be developed at the steady state with uniform concentration at the boundaries.

In the center channel of the microfluidic platform, KIC cells were uniformly implanted in 2mg/ml type I collagen mixture (Corning Inc., NY, USA) supplemented with 10X PBS, NaOH, HEPES solution, FBS, Glu, P/S, and cell-culture level distilled water. Initial cell density was 8×10^5^ cells/ml consistently for all groups. After loading, the cells in the collagen matrix were cultured with basic mediums for 24 hours. Then, cells were exposed by engineered signal environment accordingly.

### Pressure driven flow in the microfluidic platform

We controlled the low Reynolds flow through the collagen matrix (0.5–3μm/s) that corresponded to the interstitial flow rate of the tumor microenvironment. [52,53]. In controlling the flow rate inside the collagen matrix, we considered the Brinkman equation:

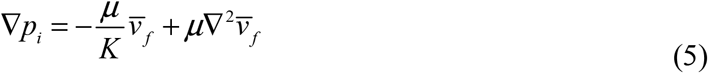

Where 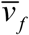 is the average flow velocity, *μ* is a dynamic viscosity, and K is the permeability of the culture medium in a type I collagen matrix of 2mg/ml. [53,73] In the literature, the permeability K in a type I collagen matrix of 2mg/ml has been reported to range from 10^−14^–10^−13^ m^2^. [74-76]. Based on that, we averaged the value range of reported permeability K, calculated as K = 5×10^−14^ m^2^. The pressure variance was applied between the source and sink channels by controlling the hydrostatic pressure levels of each reservoir, respectively. To control the flow velocity of ∼1 μm/s, we considered the pressure differences ΔP (P_source_ – P_sink_) = ∼2mmH_2_O, adapting ∇p ∼ 19.6 Pa/mm in the center channel. The hydrostatic pressure differences are controlled by applying the medium level differences between two channel reservoirs with a presence of drain flow. The drain flow was applied aiming to maintain the pressure difference between the channels consistently. Here, we assumed that the drain flow at the sink channel is not critically interrupted the interstitial flow at the center channel with relatively small area of the interface. The consistent drain flow at the sink channel (Q_drain_=10μl/h) was applied considering potential pressure drop caused by the flow. The drain flow was achieved by connecting the sink channel with syringe pump (NE-1000-ES, New Era pump system, USA).

To verify the scale of the controlled flow rate, we measured fluorescent beads’ (0.2μm diameter) trajectories. The average ± standard error of the collected particle velocities was 1.5 ± 0.048 μm/s (**Figure S1**). By using the measured value of the flow velocity, the permeability K for 2mg/ml type I collagen matrix was calculated as 8 × 10^−14^ m^2^ where μ = 0.84cP for DMEM [53], which is within comparable scale with the reported permeability range of 10^−14^–10^−13^ m^2^[74-77].

### Characterization of the directed cell migration

Live-cell time-lapse imaging with an inverted microscope (Olympus IX71, Japan) is utilized to characterize the cell migration. A stage top incubator allows maintaining the microfluidic platform at 37ºC with 5% CO_2_ condition during imaging as described in our previous studies. [47] Migrating eKIC cells were captured every 5 minutes for 3 hours. The time-lapse images are captured 3 hours after applying either chemical or pressure variances to give an adjustment time for stable environmental condition. The bright-field time lapse images are segmented to analyze cell trajectories by using ImageJ. A specific cell region is determined by the image contrasts which provides clear boundaries between cells and background. Then, cell centroids are collected in the converted monochrome images. A collection of the centroids of cell areas at different time points are defined as a cell trajectory. In collecting cell trajectories, we reject trajectories of cells under division and the stationary cells. This is because the dividing cells could affect for cell polarity [78] and the stationary cells could underestimate the cell movement characteristics. The stationary cells were defined when a cell’s total trajectories were less than the estimated cell diameter.

The directed cell migration is characterized by motility and directional accuracy. [47] The direction of the cell trajectories is analyzed based on the direction of environmental signals. We measure directional accuracy using the directional accuracy index (DAI)

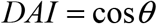

where *θ* is the angle between the net displacement of a trajectory and the environmental cue direction. A straight line connecting the initial and final points of a trajectory indicates a displacement. For the chemotaxis, the direction of the environmental signal is along the concentration gradient direction from low to high. When the interstitial flow is applied as an environmental signal, we compare the cell bias with the upstream direction of the flow along the flow streamline, considering the recent studies reporting that the cells were stimulated toward the upstream direction. [74] When both chemical gradient and interstitial flow are spontaneously applied, the reference direction of the signals is determined as the chemical gradient direction. The DAI range is between -1 and 1. DAI = 1 indicates that the cell is perfectly biased to the environmental signal direction, whereas DAI = 0 means that the cell is showing random motion. On the other hand, DAI=-1 indicates that the cell moves toward the completely opposite direction to the environmental signal. Thus, higher DAI indicates that the cell migration is accurately following the reference direction. Cells show distributed DAIs throughout the range of -1 to 1 due to the nature of cell response to the attractant. In the distribution, a median DAI represents a result from one experiment trial. More detailed description about DAI is stated in the previous studies[47]. Here, the cell path is measured from a trajectory taken every Δ*t* = 5 minutes, and total duration of the trajectories is three hours.

### Statistical analysis for experiments

All experimental controls were repeated until the number of trajectories in each case > 50 trajectories. A trajectory was evaluated with a quantified DAI and a speed. To compare the directional accuracy, the distribution of DAIs was reported in box plots with distribution of data points. A data point in the box plots indicates the metric of a cell trajectory. Median values of the distribution were statistically examined with Mann-Whitney nonparametric test where the statistical significance was evaluated when U <0.05 in **Figure 1D and Figure 3C**.

