## Supplementary Information for "Cells function as a ternary logic gate to decide migration direction under integrated chemical and fluidic cues"

**Supplementary Information – Physical model**

The dynamics of *x, y, b* and *m* from the network in **Figure 4A** are given by,

$$\dot{x} = k_1 c - k_2 x$$

$$\dot{y} = k_3 f - k_4 y$$

$$\dot{b} = (k_5 + k_6 x + k_7 y)(b_0 - b) - k_8 b$$

$$\dot{m} = k_9 b - k_{10} m$$

Where $b_0$ is the total amount of molecules of type A and B. $k_i$s are rates of reactions, *c* is the external chemical concentration and *f* is the corresponding effect due to the flow. The underlying assumption of putting *f* and *c* in the same footing is that both the chemical signal and flow activates internal molecules X and Y in the cell. The steady state expression of *m* is,

$$m = \frac{k_9}{k_{10}} \frac{\left[k_5 + \left(\frac{k_6 k_1}{k_2}\right) c + \left(\frac{k_7 k_3}{k_4}\right) f\right] b_0}{k_5 + k_8 + \left(\frac{k_6 k_1}{k_2}\right) c + \left(\frac{k_7 k_3}{k_4}\right) f}$$

To determine the difference of *m* between two halves of the cell we use,

$$\Delta m \cong \frac{\partial m}{\partial c} \Delta c + \frac{\partial m}{\partial f} \Delta f$$

Where the first term is the contribution due to the chemical gradient and the second term is the contribution due to the flow. Simple algebra leads to the expression,

$$\Delta m = \frac{k_8 k_9 b_0}{k_{10}(k_5 + k_8)^2} \frac{\left[\left(\frac{k_6 k_1}{k_2}\right) \Delta c + \left(\frac{k_7 k_3}{k_4}\right) \Delta f\right]}{\left[1 + \left(\frac{k_6 k_1}{k_2(k_5 + k_8)}\right) \bar{c} + \left(\frac{k_7 k_3}{k_4(k_5 + k_8)}\right) f\right]^2}$$

Where $\bar{c}$ is the background chemical concentration and $\Delta c$ is the difference of chemical molecules between two halves of the cell and *f* and $\Delta f$ is the corresponding effect due to the flow. $\bar{c}$ and $\Delta c$ can be rewritten in terms of known quantities as $a'g$ where $a'$ is the cell length and $g$ is the chemical gradient.

We can also define the following dimensionless parameters,

$$\eta = \frac{k_8 k_9 b_0}{k_{10}(k_5 + k_8)};\ \beta = \frac{k_6 k_1}{k_2(k_5 + k_8)}; \phi_1 = \left(\frac{k_7 k_3}{k_4(k_5 + k_8)}\right)\Delta f; \phi_2 = \left(\frac{k_7 k_3}{k_4(k_5 + k_8)}\right) f$$

To get the expression in **Eq.3** in the main text,

$$\Delta m = \eta \frac{\beta a' g + \phi_1}{(1 + \beta \bar{c} + \phi_2)^2}$$

Now observations from our own cancer cell migration experiments suggest there is always a random component to migration (mean and median DAI is always less than the maximum possible value 1). Moreover, DAI is bounded between -1 and 1 and $\Delta m$ is unbounded. So, to draw a more direct analogy of $\Delta m$ to DAI we define the probability distribution of migration angles $\theta$ as a function of $\Delta m$ as a biased random walk model (Varennes et al., 2019)

$$p(\theta) = \frac{1-\alpha}{2\pi} + \frac{\alpha e^{-(\Delta m)\cos\theta}}{2\pi I_0(\Delta m)}.$$

Where, $0 \leq \alpha \leq 1$ is the maximum possible mean DAI ($\langle \text{DAI} \rangle = \int_0^{2\pi} cos(\theta)\, p(\theta) d\theta$) and $I_0$ is the modified Bessel function of the first kind. The value of $\Delta m$ suggests how close mean DAI is to $\alpha$. The larger and positive $\Delta m$ is the closer mean DAI is to $\alpha$ and as $\Delta m \to \infty, \langle \text{DAI} \rangle \to \alpha$ .

Now we set $\alpha, \eta, \beta, \phi_1, \phi_2$ are the unknown parameters. Among these we set $\alpha$ equal to the maximum mean DAI in experiments in **Figure 4C**. The other four unknown parameters are set to give minimum total absolute error between the experimental median DAI and median DAI using our model in experiments shown in **Figure 4C**. The best set of parameter values are,

$\eta = 3003.9; \beta = 0.9073; \phi_1 = 0.0018; \phi_2 = 1.6612.$

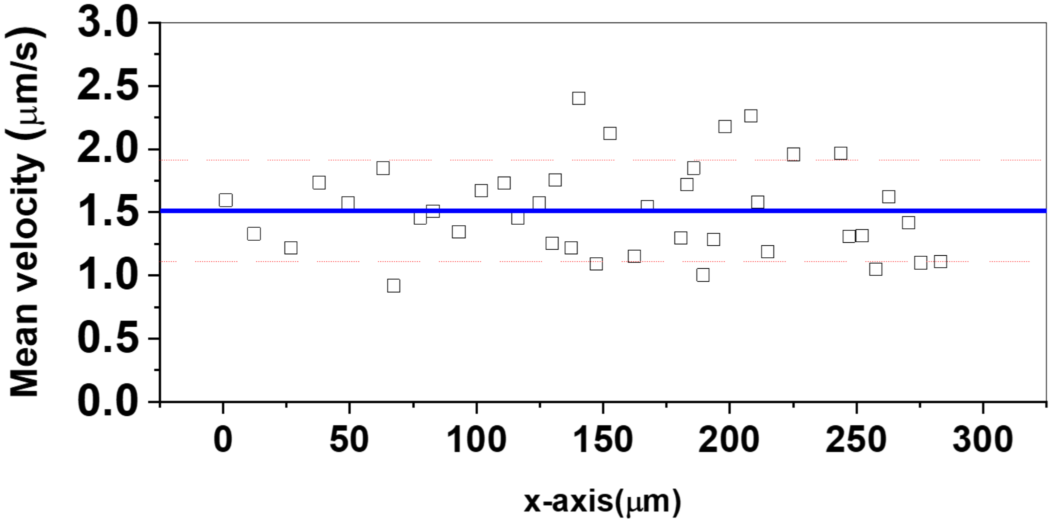


**Figure S1. Flow velocity simulated by fluorescent polystyrene beads of 0.2μm diameter.** Dot: The velocity of the collected particles distributed along with the x-locations. Blue line indicates average of all collected particles' velocity and red dash lines indicate standard error.

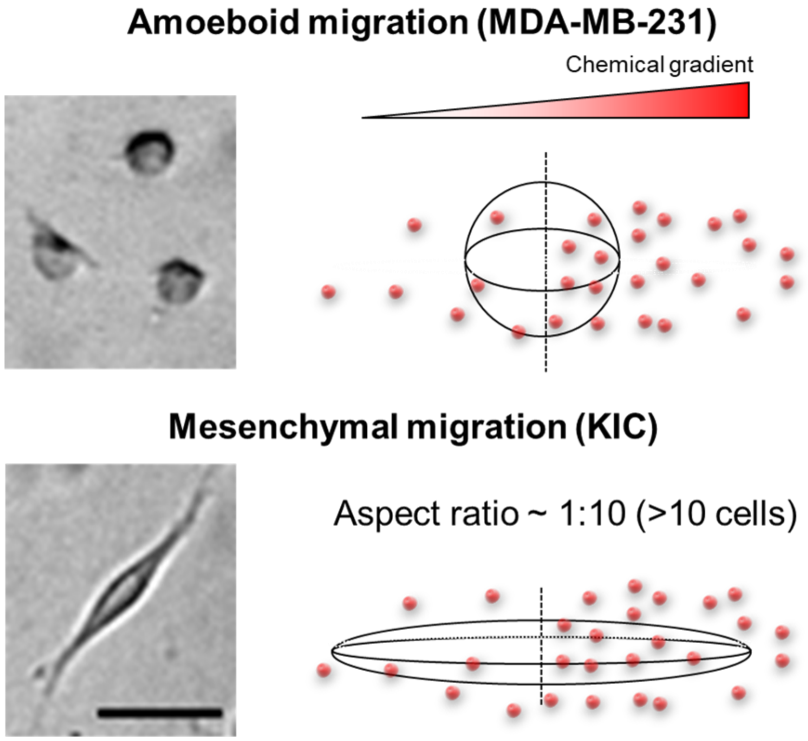


$$S = \frac{\text{Surface area of ellipsoid (mesenchymal)}}{\text{Surface area of sphere}}$$

**Figure S2. Shape factor to adjust the mesenchymal morphology of KIC cells in relative gradient of the chemical concentration across the cell body ( $\gamma$ ) .**

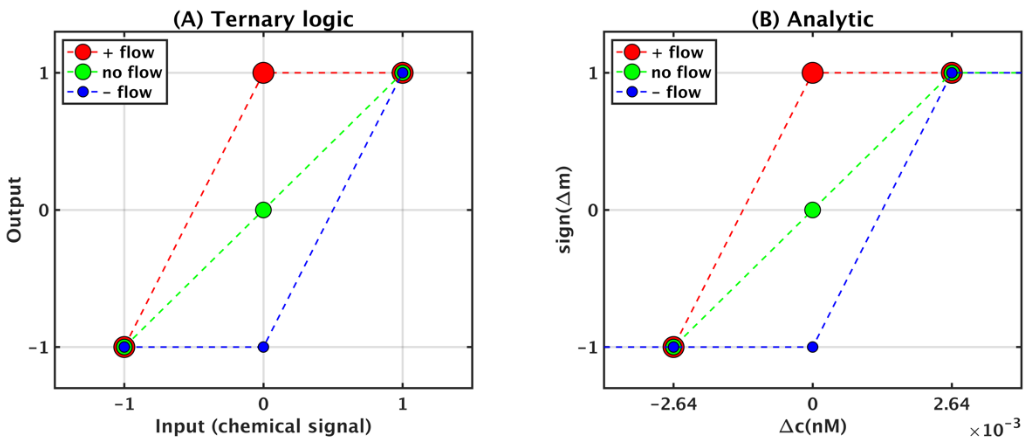


**Figure S3. Relationship between ternary logic gate model and analytic expression for $\Delta m$**

There is a one-to-one correspondence between the (A) ternary logic gate model as shown in **Figure 5** analytic expression for $\Delta m$ in **Eq. 3**. The expression of the output ($Y$) in **Figure 5** can be written as $Y = (f(1-|g|)+g) \times (1-\bar{c})$, where $g = 1$ if $\Delta TGF$ is positive and above the detection limit, $g = 0$ if $\Delta TGF$ is below the detection limit or absent, and $g = -1$ if $\Delta TGF$ is negative and above the detection limit;; $f = 1$ if the pressure gradient is positive (such that the flow is in the negative direction and cells would move in the positive direction upstream), $f = 0$ if flow is absent, and $f = -1$ if the pressure gradient is negative; and $\bar{c} = 1$ if the background TGF concentration is above the saturation limit (~10 nM as indicated in **Figure 4D**), and $\bar{c} = 0$ if the background TGF concentration is below the saturation limit.. In A, the vertical axis is $Y$, the horizontal axis is $g$, the legend indicates $f$, and $\bar{c} = 0$. In B, the sign of Eq. 3 is plotted with the parameters fitted in the main text, where $2.64 \times 10^{-3}$ nM is the concentration at which $\Delta m$ changes sign in the presence of flow. In both cases, we see that the output only reflects the flow state if the chemical gradient is absent (A) or sufficiently small (B).

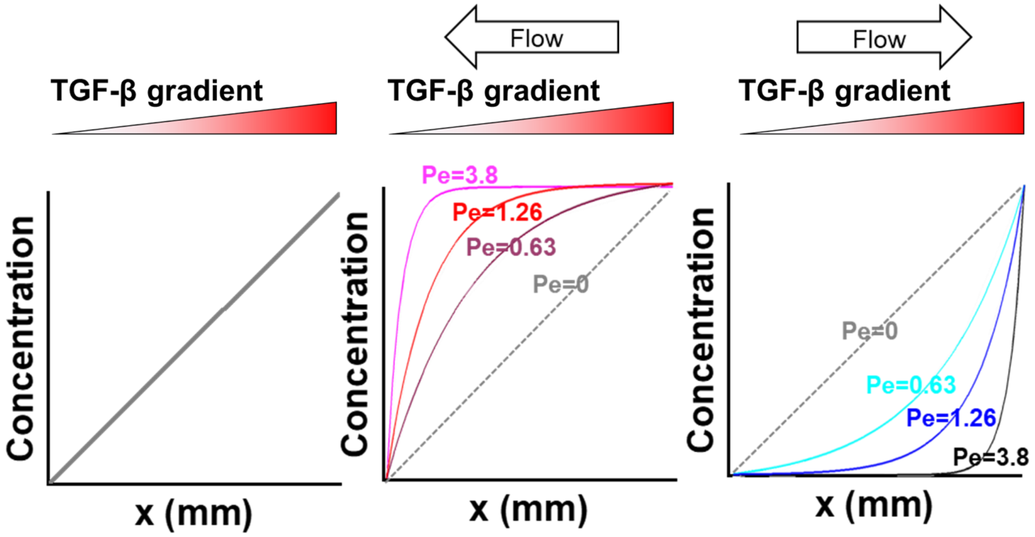


**Figure S4. Concentration profiles of the chemical cue integrated with fluidic cue with various Pe numbers (Pe=0, 0.63, 1.26, and 3.8) for each flow condition of (A) no-flow, (B) parallel flow, and (C) counter flow.**
